# Changes to task relevance in redundant motor environments distinguish structure learning from adaptation

**DOI:** 10.64898/2026.10.26.762093

**Authors:** Pavithra Rajeswaran, Leo R. Scholl, Lydia I. Smith, Amy L. Orsborn

## Abstract

Motor actions often exceed the dimensionality of task feedback, which creates control redundancy that requires learners to identify which movement dimensions influence task outcomes. The computations that shape learning in redundant tasks, especially when task-relevant spaces change, remain unclear. We studied this by combining established visuomotor perturbation approaches with a redundant motor interface reaching task. Two monkeys learned a 3D hand position to 2D cursor position mapping that allowed us to define and manipulate the task-relevant space. After animals learned an initial task-relevant space, we introduced two types of perturbations: Relevance Preserving (RP), which rotated hand-cursor relationships within the learned task relevant space, and Relevance Altering (RA), which altered the task relevant space. We hypothesized that RP perturbations would engage error-based adaptation (observed in non-redundant tasks), whereas RA perturbations would engage additional learning mechanisms associated with acquiring new skills. RP perturbations were learned rapidly during training (online) and generalized to the initial learned mapping, consistent with error-based adaptation. In contrast, adapting to the first RA perturbation took longer, engaged learning outside of training (offline), and did not generalize to the initial learned mapping. Animals also adapted faster across repeated exposures to new RA mappings and changed how they generated movements to learn each new mapping, consistent with computations associated with structure learning. These findings suggest that changing task relevance shifts motor learning from adaptation toward structure learning, which has implications for models of motor learning and sensorimotor therapies like brain-computer interfaces.

**Significance Statement:** Most real-world motor tasks provide many ways to achieve a goal. This redundancy requires learning the task’s structure, including which actions are relevant, but many studies do not incorporate redundancy. We added redundancy to a visuomotor perturbation paradigm to selectively perturb task-relevant spaces. Perturbed visual-motor mappings that preserved task-relevant spaces produced rapid adaptation, while mappings that altered task-relevant spaces yielded signatures of structure learning, including history-dependent meta-learning. Our study reveals a continuum between forms of learning typically studied separately and demonstrates that cognitive inferences about task structure shape exploration in redundant motor tasks. Our work highlights the need to consider structure learning in therapies like brain-computer interfaces, and advances our ability to study the computations involved in real-world motor skills.

## Introduction

Movements typically involve more degrees of freedom than required to achieve a goal. This *redundancy* takes two forms: s*olution redundancy*, where multiple movements (trajectories, postures) can achieve the same outcome (Bernstein, 1967; Yang et al., 2007), and c*ontrol redundancy,* where the dimensionality of movement exceeds that of task feedback (So et al., 2012; Law et al., 2014). For instance, using an arm to control a 2D cursor pairs a high-dimensional action space with 2-dimensional feedback. Solution redundancy has been more widely studied than control redundancy. Learning motor skills in control-redundant environments requires identifying which actions influence task outcomes – the *task relevant* subspace. How task relevant subspaces are initially learned, and how learning unfolds when task relevance changes, are largely open questions.

Motor learning tasks often restrict control redundancy by aligning movement and feedback spaces (Shadmehr and Mussa-Ivaldi, 1994; Krakauer, 2009; Krakauer et al., 2019). Common paradigms like a 2D cursor controlled by a 2D manipulandum ensure that all movements produce task-relevant feedback, unambiguously relating task errors to actions. These paradigms enabled major advances in models of error- and internal-model-based adaptation (Wolpert and Flanagan, 2001; Shadmehr et al., 2010).

However, non-control-redundant tasks cannot probe computations that are central to learning motor skills like identifying task-relevant actions.

Redundant behavioral tasks have been used to study how task-relevant spaces are identified, with a particular focus on the role of movement variability. Redundancy partitions movement into portions that do and don’t influence task performance (*task-relevant* and *task-irrelevant*, respectively) (Scholz and Schöner, 1999; Müller and Sternad, 2004). Most work on solution redundancy characterized the structure of movement variability after a task-relevant subspace is learned (Scholz and Schöner, 1999; Todorov and Jordan, 2002; Scott, 2004). Complementary work on how task-relevant subspaces are acquired suggests that action-outcome relationships are inferred by actively regulating variability based on knowledge of task structure (Wu et al., 2014; Pekny et al., 2015; Therrien et al., 2016; Dhawale et al., 2017). These forms of skill acquisition have been examined behaviorally in humans (Dal’Bello and Izawa, 2021; Kobayashi and Nozaki, 2024) and its neural mechanisms have been well-characterized in songbirds (Fee and Goldberg, 2011; Dhawale et al., 2017).

If knowledge of task structure contributes to initially learning a task relevant space, does it also inform how we *adapt* to new tasks? Adapting to a new task-relevant space could engage error-based adaptation (observed in non-redundant tasks), or structure learning, or interactions of both. Addressing this question requires manipulating the task-relevant space without altering task goals. Most redundant task paradigms, however, confound task relevance with task goals or other factors. Myoelectric and brain computer interfaces, for example, involve control redundancy and probe learning without modifying task goals, but neurophysiological factors such as muscle synergies and synaptic connections may constrain the space of inputs (actions) someone can generate (d’Avella and Pai, 2010; Berger et al., 2013; Sadtler et al., 2014; Oby et al., 2019). These partially unknown constraints make it difficult to isolate learning related to task-relevance.

To isolate learning computations, we designed a sensorimotor task with control redundancy but minimal biological constraints. Monkeys used 3D hand movements to control 2D cursor movements. After learning a “baseline” mapping, we introduced perturbations that either preserved the learned task-relevant subspace (relevance preserving, RP) or defined a new subspace that included movement in previously task-irrelevant dimensions (relevance altering, RA). We hypothesized that these two perturbations would engage distinct learning mechanisms: RP perturbations would be solved via error-based adaptation, whereas RA perturbations would engage additional learning mechanisms. We observed clear distinctions in the timescales of learning and generalization patterns between these two perturbations that were consistent with this hypothesis. RA perturbations also showed a unique dependence on prior training history and changes in how animals generated movement variability, suggesting that structure learning contributed to adapting to new task spaces. Together, these findings demonstrate that task-relevant spaces can influence the learning mechanisms employed to shape motor behaviors.

## Materials and Methods

### Animal selection and surgical procedures

All animal procedures were approved by the IACUC at the University of Washington. Two male rhesus macaques (*Macaca mulatta*) were selected as experimental subjects. Monkey A (10 years old, 10.6 kg) and Monkey B (9 years old, 11.3 kg) were pair-housed at the Washington National Biomedical Research Center with a 12h light/12h dark cycle. Both monkeys had scheduled access to water when on the study. Prior to data collection, animals underwent surgery to implant a titanium head restraint.

### Experimental Design

Monkeys were trained to perform arm movements to control the movements of a cursor displayed on a two-dimensional display (Fig. 1A). Cursor control and visual stimuli were implemented using BMI3D (http://github.com/aolabNeuro/brain-python-interface). The display (16:9 LCD, 21.5 cm vertical size) was positioned 28–30 cm from the animal’s eyes, and monkeys moved their right arm freely just below the screen within a 20 × 20 × 20 cm motion capture volume. Visibility of their right hand was limited by a polycarbonate sheet placed horizontally between the display and the movement volume. The monkeys’ left arm and head movements were restrained. The position of the cursor was determined by a linear projection of the monkey’s right hand position (see <u>Hand to Cursor mapping</u>). We used a marker-based optical tracking system (NaturalPoint, Inc., Corvallis, OR) to capture the three-dimensional position of the animal’s hand in real time at a rate of 240 Hz. Monkeys wore a lightweight glove equipped with infrared-reflective markers to enable motion tracking. All data streams, including hand position, cursor position, and task events, were temporally aligned using synchronized clocks via a data-acquisition system (eCube, White Matter Research, Montreal CA).

**Figure 1:**
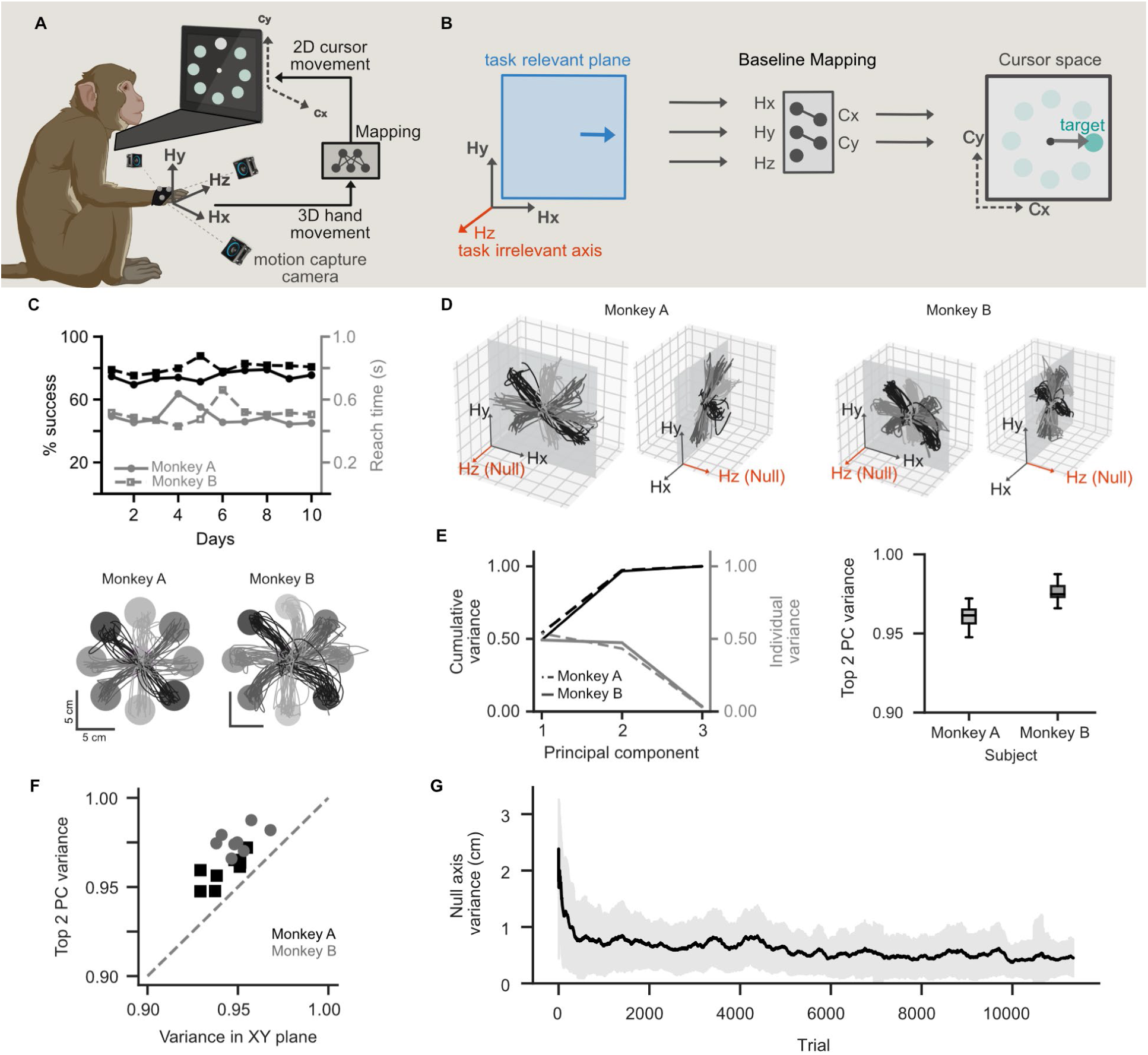
Monkeys learned a planar solution to baseline 3D hand to 2D cursor mapping. **A)** Experiment schematic. Monkeys moved their unseen right arm in 3D space to control a 2D target acquisition task displayed on a computer monitor. 3D hand movements were tracked in real-time with motion capture cameras ([Hx, Hy, Hz]), which were mapped to 2D cursor movements ([Cx, Cy]) via an experimenter-defined mapping. **B)** Schematic illustrating the relationship mapping (grey box, middle) between hand movement space (left) and cursor movement space (right) for the baseline mapping. Hx, Hy were mapped to Cx, Cy respectively, while Hz was task irrelevant, which defined a 2D plane of movement space that is task-relevant (blue). **C)** Top: Success rate (black) and reach time (gray) across days for Monkeys A (circles) and B (squares). Bottom: Example cursor trajectories to each of the eight targets, color coded by direction (left: Monkey A; right: Monkey B). Scale bars denote 5 cm. Data chosen from 10 consecutive baseline days randomly selected before perturbations were introduced. **D)** Example 3D hand trajectories corresponding to cursor trajectories shown in (C) (left: Monkey A; right: Monkey B). All views include the task relevant plane (XY plane; grey) and the task-irrelevant direction of movement (Hz axis; orange). **E)** Left: Cumulative variance (black) and variance explained by each PC (gray) for hand movements on one example day (first of the 10 days shown in C), for Monkey A (dashed lines) and Monkey B (solid lines). Right: Cumulative variance explained by top 2 PCs for each monkey, across 10 baseline days before perturbations were introduced (same data as in C). Boxes indicate inter-quartile range, lines indicate medians, whiskers extend to 1.5x IQR.**F)** Proportion of hand movement variance explained by the top 2 PCs vs hand movement variance captured in the baseline (XY) task plane (see methods) for Monkey A (black, squares) and Monkey B (grey, circles). Each datapoint reflects data from a single day. Gray dotted line indicates unity line. **G)**Task irrelevant variance across trials during baseline training for Monkey B. Solid line indicates rolling mean and shaded region indicates rolling standard deviation computed from a rolling window of 21 trials.

#### Point to point reaching task

Monkeys performed a delayed point-to-point reaching task. Each trial was self-initiated by holding the cursor inside a central target for 200 ms. Then, one of eight peripheral targets, equally spaced at 45° intervals appeared 6.5 cm from the center of the screen. The task required the cursor to remain inside the center target during a variable delay of 100–600 ms (uniformly distributed) until the center target disappeared, which cued the monkey to initiate a reach. Reaches to the peripheral target did not have any experimentally imposed time constraints, and rewards were delivered for successful reaches only after holding within the peripheral target for 200 ms. After a trial ended, with or without reward, animals returned to the center to initiate the next trial after a randomized waiting period (Monkey A: 0.04–0.06 s or 0.12–0.3 s; Monkey B: 0.2–0.5 s).

#### Hand to Cursor mapping

Our experimental setup created redundancy between the hand and task spaces, which we used to investigate how task-relevance influences learning. The task-relevant space of actions was defined by the linear mapping (*M*) from 3D hand position space (*H*) to 2D cursor position space (*C*) :

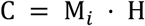

where:

∈ ℝ^3^ *^x^* ^1^is the hand position vector denoting X, Y and Z movement axis,
*C* ∈ ℝ^2^ *^x^* ^1^is the cursor position vector denoting X and Y axis, and
M*_i_* ∈ ℝ^2^ *^x^* ^3^ is the linear mapping matrix for condition *i*.

Because there are more input dimensions (3D) than output dimensions (2D), the mapping introduces redundancy. Specifically, the mapping defines a 2D task-relevant *plane*, in which movement affects the cursor, and a task-irrelevant 1D *axis* along which movements do not affect the cursor.

Monkeys first learned a mapping which we define as baseline, *M_base_* . This mapping defined cursor space to be a 1:1 linear projection of the horizontal (*X*) and vertical (*Y*) components of hand movement:

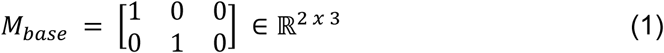

This baseline mapping defines the *XY* hand movement plane as the task relevant plane. Movements along the depth (*Z*) axis (i.e., the third column of *H*) have no contribution to cursor movement. We subsequently perturbed M to either preserve or alter the task relevant plane.

##### Task-relevance preserving perturbations

We applied a second linear transformation (*R*) to the baseline mapping (*M_base_*) to create mappings that altered hand-cursor relationships while preserving the task-relevant space (RP mappings, *M_rp_*). For our study’s baseline mapping *(eq. 1)*, RP mappings maintained *XY* as the task-relevant plane while changing how the cursor moved in response to hand movements within the *XY* plane (Fig. 2A). In the space of possible transformations, we studied RP perturbations that applied a fixed rotation angle θ about the *Z* axis ranging from 20° to 90° (±20°, ±45°, ±60°, ±90°). This corresponds to multiplying the baseline mapping by a two-dimensional rotation matrix *R* ∈ ℝ^2*x*2^ :

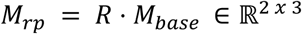

**Figure 2:**
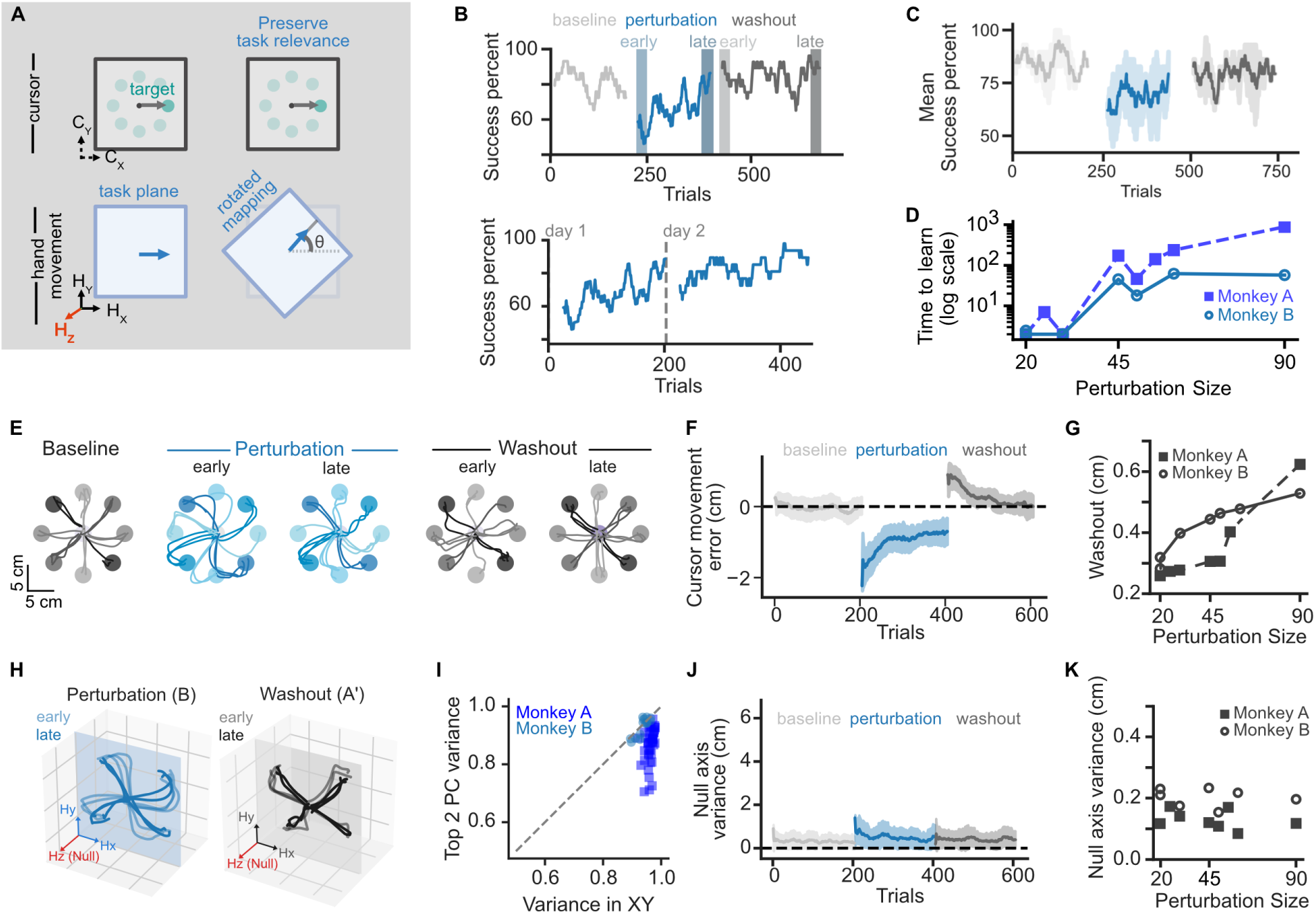
Relevance preserving perturbations learned within day and show generalization. **A)** Perturbation schematic. Cursor movement space (top) and hand movement space (bottom) for the baseline mapping (left) and a relevance-preserving perturbation (right). Relevance preserving perturbations rotated the mapping by an angle (Ɵ) within the baseline task relevant plane (blue arrow) while keeping the task relevant plane and the task irrelevant axis unchanged. **B)**Top: Success percent vs trials during baseline (A, light grey), perturbation (B, blue) and washout (A’, dark grey) blocks on day 1 for example 45° perturbation from Monkey B. Shaded vertical regions mark the early (light blue in B, light grey in A’) and late (dark blue in B, dark grey in A’) trial blocks. Bottom: Success percent during the perturbation block across two consecutive days of training within a learning series (same series shown in the top panel). Vertical grey line indicates boundary between days. **C)** Mean success percent across trials in the 45° RP mapping from Monkey B (N = 2 days). Colors denote baseline (light grey), perturbation (blue) and washout (dark grey) blocks, as in (B). Solid lines indicate mean rolling success percent (sliding window 21 trials) and shaded region indicate standard deviation of rolling mean. **D)** Time to learn as a function of absolute perturbation size for Monkey A (dark blue, squares) and Monkey B (light blue, circles). Each point is one RP learning series (N = 15 series across two monkeys). **E)** Example successful cursor trajectories (color coded by target) during baseline (gray), early and late perturbation (blue; left and right, respectively), early and late washout blocks (gray; left and right, respectively) from Monkey B. Data are shown from day 1 of exposure to a 45° perturbation. **F)** Cursor movement error (deviation from a straight line) across trials in the 45° RP mapping from Monkey B (N = 2 days). Format is the same as panel (C). **G)** Washout size (see methods) as a function of absolute perturbation size for Monkey A (grey, filled squares) and Monkey B (grey, unfilled circles). Each point is one RP learning series (N = 15 series across two monkeys). **H)** Hand movement trajectories in 3D during perturbation (blue, left) and washout blocks (gray, right) for Monkey B. Color shading denotes trial order, with early trials shown in lighter colors. Only four target directions are shown for clarity. Blue and grey planes show the task relevant plane in respective blocks. **I)** Proportion of hand movement variance explained by the top 2 PCs vs hand movement variance captured in the baseline (XY) task plane (see methods) for Monkey A (dark blue, squares) and Monkey B (light blue, circles). Each datapoint reflects data from a single day. Gray dotted line indicates unity line. For C, F and I, solid line shows the rolling mean and shaded area indicates rolling standard deviation. **J)** Null-axis variance across trials in the 45° RP mapping from Monkey B (N = 2 days). Format is the same as panels (C) and (F). **K)** Null-axis variance during the first 60 trials of the washout (A′) block vs. perturbation size for Monkey A (grey, filled squares) and Monkey B (grey, unfilled circles). Each point is one RP learning series (N = 15 series across two monkeys).

We use rotation angle as a measure of perturbation size since it captures the deviation – within the original task plane – from the baseline mapping. Importantly, because the perturbed task plane remained fully aligned with the learned baseline plane, movements in the baseline plane projected entirely to the perturbed plane and thus generated task feedback (i.e., cursor movement).

#### Task-relevance altering perturbations

We applied a second linear transformation (*QQ*) to the baseline mapping (*M_base_*) to create mappings that altered hand-cursor relationships *and* altered the task-relevant space (RA mappings, *M_ra_*). This manipulation is enabled by our redundant mapping paradigm. For our study’s baseline mapping (eqn. 1), RA mappings defined a task-relevant plane that was distinct from the *XY* plane and may also alter hand-cursor relationships within the reference frame of the new task-relevant plane (Fig. 3A). This corresponds to post-multiplying the baseline mapping by a 3D rotational matrix *QQ* ∈ ℝ^3^ *^x^* ^3^ :

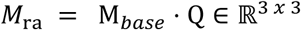

**Figure 3:**
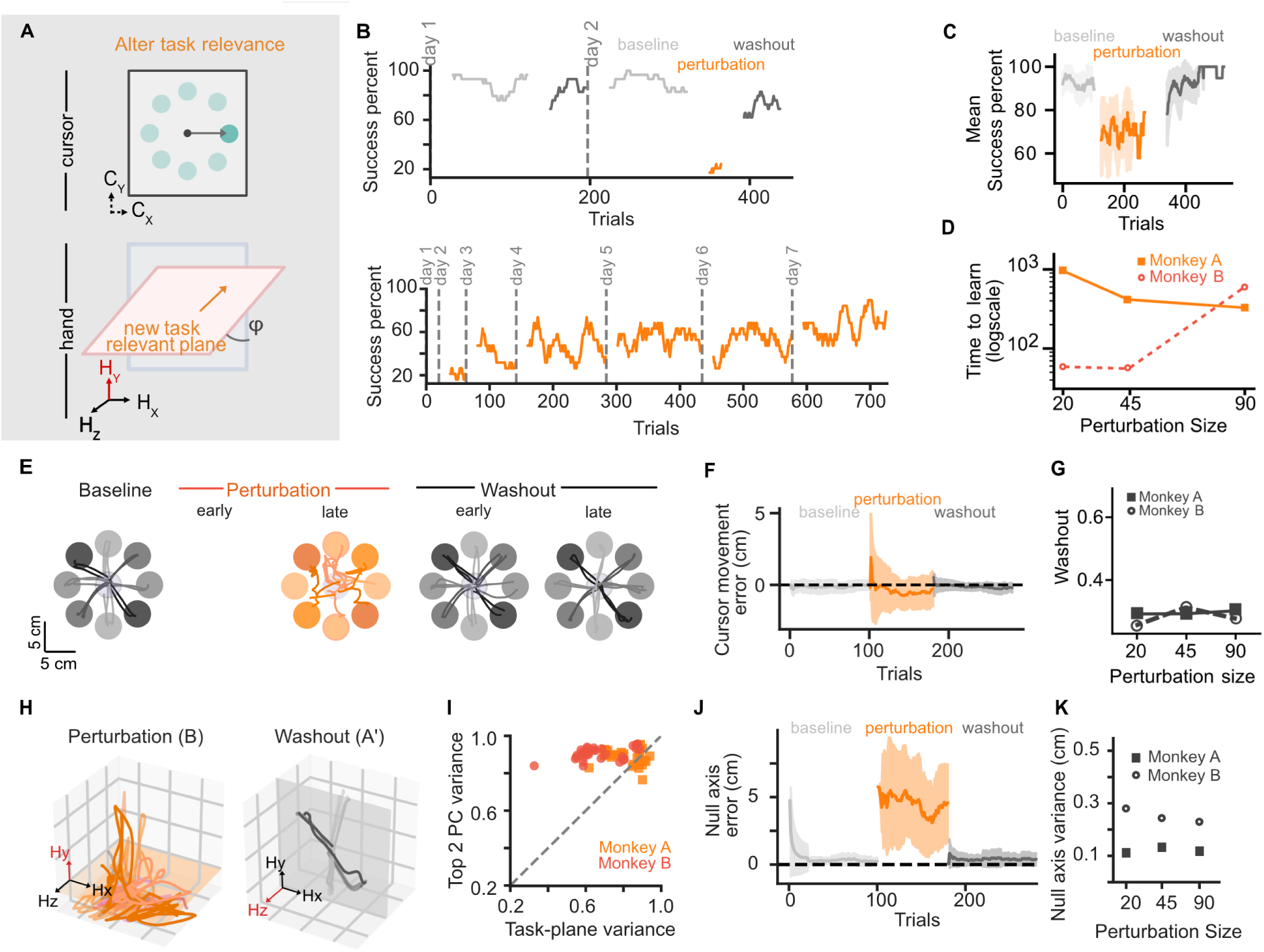
Relevance altering perturbations required multi-day learning and showed no generalization. **A)** Perturbation schematic. Relevance altering perturbations rotated hand position by an angle φ about the X axis (orange), thus changing the task-relevant plane to something different from the baseline mapping (blue). **B)** (Top) Success percent across trials during baseline (A, light gray), perturbation (B, orange), and washout (A’, dark gray) blocks on day 1 and 2 for an example 90° perturbation from Monkey B. The number of successful trials completed on day 1 was fewer than the size of the rolling window. (Bottom) Success percent during B block across multiple days of the same learning series from Monkey B. Vertical grey lines indicate boundaries between days. **C)** Mean success percent pooled across all days from 90° RA mapping from Monkey B (N = 8 days). Colors denote baseline (light grey), perturbation (orange) and washout (dark grey) blocks, as in (B). Solid lines indicate mean rolling success percent (sliding window 21 trials) and shaded region indicate standard deviation of rolling mean. **D)** Time to learn vs. perturbation size for Monkey A (light orange, squares) and Monkey B (dark orange, circles). Each point is one RA learning series (N = 6 series across two monkeys). **E)** Example successful cursor trajectories to eight targets (color coded by target) during baseline (grey), early and late perturbation (orange; left and right, respectively), and early and late washout (grey; left and right, respectively) from Monkey B. Data are shown from day 2 of 90° perturbation. No successful reaches in early perturbation block. **F)** Cursor movement error (deviation from a straight line) across trials during baseline (light grey), perturbation (orange) and washout (dark grey) blocks from 90° perturbation from Monkey B (N = 8 days). Format is the same as panel (C). **G)** Washout size (see Methods) vs. perturbation size for Monkey A (grey, filled squares) and Monkey B (grey, unfilled circles). **H)** 3D hand trajectories during perturbation (orange, left) and washout blocks (grey, right) for Monkey B. Each color represents a target direction. Shading denotes trial order, with early trials shown in lighter colors. Only four target directions are shown for clarity. Orange and grey planes show the task relevant plane in respective blocks. **I)** Proportion of hand movement variance explained by the top 2 PCs vs hand movement variance captured in the task plane (see methods) for Monkey A (dark orange, circles) and Monkey B (light orange, squares). Each datapoint reflects data from a single day. Grey dotted line indicates unity line. **J)** Null-axis variance across trials in the 90° RA mapping from Monkey B (N = 8 days). Format is the same as panels (C) and (F). **K)** Null-axis variance during the first 60 trials of the washout (A′) block vs. perturbation size for Monkey A (grey, filled squares) and Monkey B (grey, unfilled circles).

This transformation rotated the baseline mapping in 3D hand space, thereby changing which hand axes projected into the 2D task plane. We hypothesized that these perturbations would require exploration beyond the movements monkeys made in the baseline mapping. We studied three RA mappings with varying magnitudes (±20°, ±45°, and ±90°). For ±90° RA perturbations, magnitude defined the degrees of rotation applied about the X-axis. For ±20° and ±45° RA perturbations, magnitude defined a compound rotation where the perturbation was first applied about the X-axis and subsequently about the Y-axis.

#### Training paradigm

After monkeys were trained on the baseline mapping, we studied how they learned different perturbed mappings. Each perturbation was introduced as a *learning series* across consecutive days. Table 1 provides a summary of the perturbations experienced by each monkey in chronological order. At the start of each learning series, a mapping was randomly selected from the predefined set of rotations. Each learning series was composed of daily sessions with a consistent block structure in which the animal experienced the baseline mapping (*A*), a perturbed mapping (B), and then a return to the baseline (A’). A blocks lasted a minimum of 50 successful trials (mean ± s.d.; Monkey A: 119 ± 100 trials; Monkey B: 141 ± 73 trials). The number of trials in the B perturbation block was chosen each day by the experimenter, balancing the desire to maximize exposure to the perturbation with the need to collect sufficient A′ block data given variations in animal motivation (mean ± s.d.; Monkey A: 155 ± 75 trials; Monkey B: 156 ± 79 trials). A′ blocks were collected with a target minimum of 60 trials to reliably quantify washout, though the actual number varied with animal motivation (mean ± s.d.; Monkey A: 151 ± 79 trials; Monkey B: 137 ± 89 trials). A learning series concluded when performance in the perturbation block reached at least 80% success.

**Table 1:** Training summary showing the order of perturbation mappings experienced by each monkey. RP: relevance preserving, RA relevance altering.

| Monkey A |  | Monkey B |  |
| --- | --- | --- | --- |
| Perturbation | Angle | Perturbation | Angle |
| RP | 45 | RP | 20 |
| RP | 50 | RP | 30 |
| RP | 55 | RP | 50 |
| RP | 60 | RP | 45 |
| RP | 20 | RA | -90 |
| RA | -20 | RP | 90 |
| RP | 25 | RA | 90 |
| RA | -45 | RA | 45 |
| RP | -90 | RP | -20 |
| RA | 20 | RA | -20 |
| RA | 90 | RP | -60 |
| RP | -60 |  |  |

### Data Analysis

Hand movement data and cursor positions were low-pass filtered with a 15 Hz cutoff using a zero-phase Butterworth filter and down-sampled to 100 Hz. These preprocessed data were used in all subsequent analyses of movement kinematics and task-related behavior. We visualized the temporal evolution of behavioral metrics by applying a rolling window centered on each trial and extending 10 trials forwards and backwards (total window size 21 trials), padding the start and end of data to account for edge effects. All quantifications and statistical analyses of performance changes over time or between conditions were performed using ‘early’ and ‘late’ phases of the B and A’ blocks, defined as the first 20 and last 20 trials of each block, respectively. For the A block, we used the mean across all trials to assess baseline performance.

Trial initiation imposed no constraint on the starting position along task irrelevant axis. Animals moved freely in the task irrelevant axis and started at different positions (for example, Baseline mean starting position along the Z axis for Monkey A: -1.03 ± 3.39 cm; Monkey B: -2.63 ± 2.26 cm). To aid visualizations, we re-centered each reach segment by subtracting each trial’s start position in the task irrelevant dimension (for example, we subtracted the start position in the Z axis for baseline and RP). This re-centering does not change the learning-based metrics used for our analyses, which rely on task-space position or variance calculations invariant to coordinate system redefinitions.

#### Task performance

We quantified behavioral performance using task success percentage and reach times (Fig. 1C). Success percentage was calculated as the proportion of initiated trials that resulted in a reward. We note that this does not consider trials where the monkeys failed to successfully hold at the center target. We assume these reflect errors that are less relevant to learning the hand-cursor mapping (e.g., failing to withhold a reach for the delay period). Reach time was defined as the duration (seconds) between the cursor exiting the center target and entering the peripheral target.

#### Time to learn

We compared the rate of learning across different series by examining the number of trials required to consistently achieve a certain level of task performance, which we define as *time to learn* (Fig. 2D, 3D, 5A). Time to learn was calculated using the rolling success rate, aggregating trials across perturbation blocks from all days in a series. We then computed the trial at which the success rate remained at or above 80% for more than 20 consecutive trials.

#### Within session vs Offline performance changes

Within session changes in performance during the perturbation block were computed as the change from the initial to the peak rolling success percent (Fig. 4D, F). Offline performance changes during the perturbation block were computed as the difference between the mean success at the start of a session (first 20 trials) and the mean success (across all trials) of the preceding session (Fig. 4E, F, H, I).

**Figure 4:**
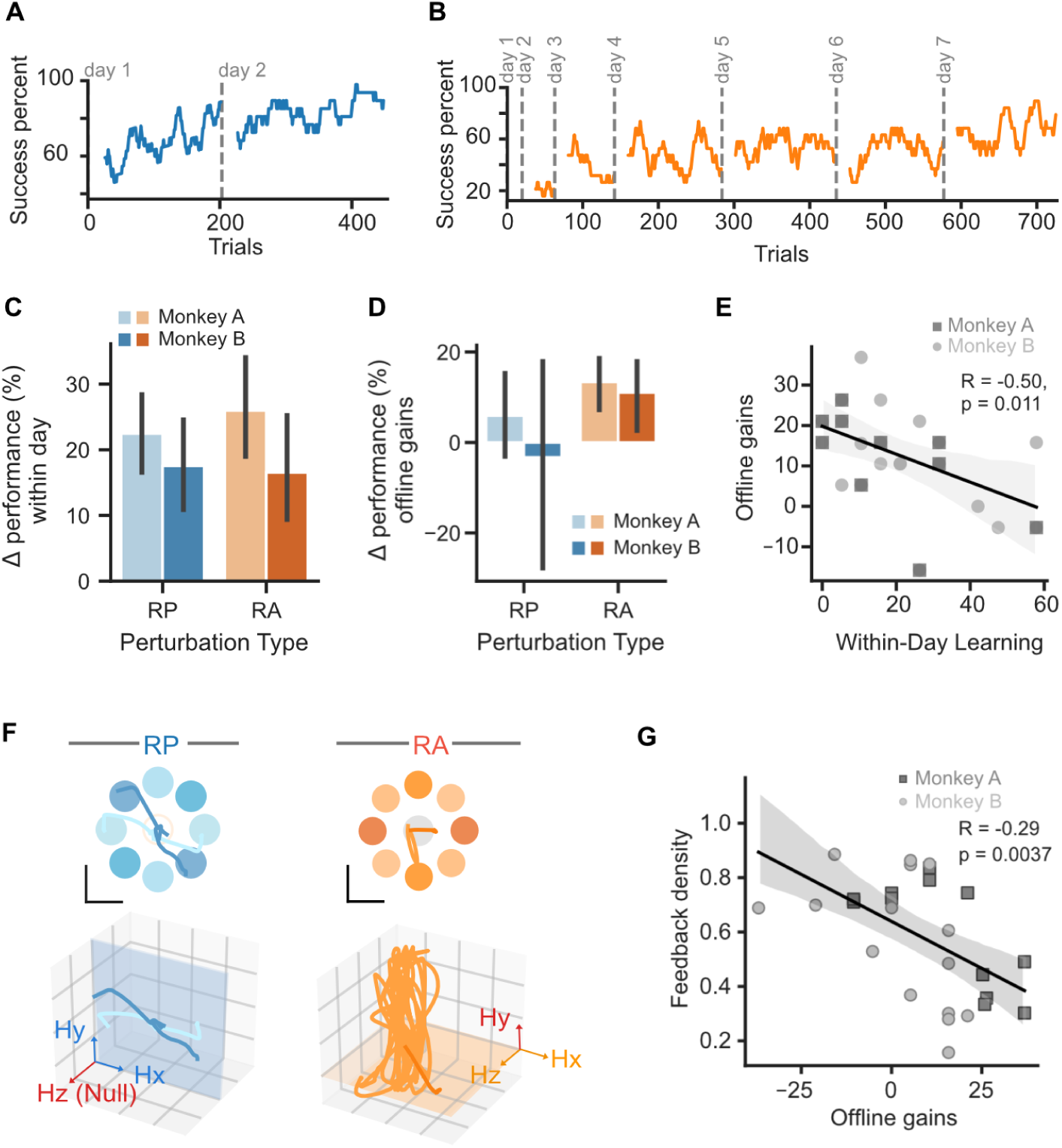
Distinct timescales of learning in relevance preserving vs relevance altering perturbations. **A)** Success percent across trials during a 45° relevance-preserving (RP) perturbation for Monkey B across two consecutive days (same as Fig. 2B, bottom). **B)** Success percent across trials during a 90° relevance-altering (RA) perturbation for Monkey B across multiple days. Same as Fig. 3B (bottom). **C)** Change in performance within a day for RP (blue) and RA (orange) perturbations, shown separately for each subject. Monkey A: 26 RP sessions (8 mappings) and 15 RA sessions (3 mappings); Monkey B: 15 RP sessions (7 mappings) and 14 RA sessions (4 mappings). Bar heights indicate the mean across sessions and error bars the bootstrapped 95% confidence interval. **D)** Offline performance gains (change in performance between days, see Methods) for RP (blue) and RA (orange) perturbations, shown separately for each subject. Offline gain was computed for each consecutive pair of days as the success percent at the start of the following day, minus the mean success percent of the preceding day. Monkey A: 16 RP day pairs (5 mappings) and 11 RA day pairs (3 mappings); Monkey B: 8 RP day pairs (4 mappings) and 10 RA day pairs (4 mappings). Format as in panel (C). **E)** Offline (across-day) gain from day N to day N+1 plotted against the within-day change in performance on day N, for Monkey A (dark grey, squares) and Monkey B (light grey, circles). Line shows least-squares fit across both subjects (r = –0.50, two-sided p = 0.011 (*), Monkey A, squares, n = 11; Monkey B, circles, n = 10; N = 21 day-to-day transitions from 7 mapping series). **F)** Example cursor trajectories (top) and corresponding 3D hand trajectories (bottom) during early learning in RP (blue, left) and RA (orange, right) perturbations. Shaded regions in the 3D plots indicate the task-relevant plane. **G)** Early feedback density on day N plotted against the offline gain measured from day N to day N + 1, pooled across animals, Monkey A, squares, n = 12, Monkey B, circles, n = 15, N = 27 sessions. Line shows the least-squares fit to all points with a 95% confidence band; Pearson *R* = −0.54, two-sided *p* = 0.0037 (**).

#### Initial learning of the task-relevant plane

We quantified how animals learned the task-relevant plane by analyzing variance in the task-irrelevant axis of hand kinematics, which corresponds to the ***Z***-axis for our baseline mapping. We computed this variance during the reach period for each trial. We quantified the timescale of variance changes by fitting the trial-by-trial variance data *V*(*t*) to an exponential decay function:

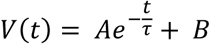

where *A* is the initial amplitude of the decay, *τ* is the decay constant, and *B* is the steady-state null axis variance. Model fits were obtained with the curve fit routine in *scipy.optimize*. The decay constant was converted to a half life 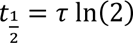, which provided the number of trials required for variance to fall halfway from its initial level toward its asymptote. Half-life values were estimated using all training sessions in which the baseline hand to cursor mapping was used, though task parameters varied across training. Initially, monkeys were trained to make a reach to a single target positioned randomly on the screen without any delay. Additional task components (initial center target, hold and delay requirements) were introduced gradually and adjusted to their final parameter settings.

#### Quantifying washout

We used task movement errors (i.e., the cursor trajectories) to quantify learning within the perturbation block and how that generalized upon return to the baseline mapping (the *washout* block). Cursor error was defined as the mean deviation from a straight line connecting the start position at the center to the target (Orsborn et al., 2012; Patton et al., 2013). We restricted our analyses to the time-period during the reach, defined as the interval between the go cue and reaching the peripheral target. We summarized the cursor error for a given trial by calculating the mean deviation across all time points during the reach. To enable cross-series comparisons (Fig. 2E & 3E), we used the mean cursor error over the first 60 trials of the A′ block as a measure of washout.

We also examined generalization patterns for hand movements using a similar approach but applied to the task-irrelevant dimension of movement. We thus computed Z-axis variance (the task irrelevant dimension for our baseline mapping) during the reach period for each trial.

#### Task-relevant variability analyses

We analyzed how 3D hand movements related to the task-relevant space by calculating the ratio of task-relevant variance (2D) to total hand variance (3D). For the baseline mapping and all RP perturbations, this corresponds to:

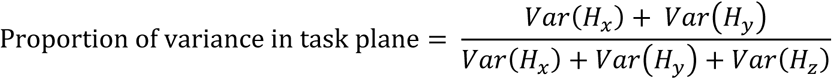

We computed this ratio during the reach segment (between the go cue and reaching the peripheral target) for each trial. Values close to 1 indicate that most hand variability lay within the task plane, whereas values close to 0 indicate predominantly task irrelevant movements. For relevance altering (RA) perturbations, where the task relevant plane was rotated, we used the perturbation-specific mapping matrix to calculate task-relevant variance.

We also characterized the distribution of variance in hand movements in 3D space independent of spatial coordinates using Principal Component Analysis (PCA). PCA removes differences in coordinate system, allowing us to compare hand movements across blocks with different mappings. We performed PCA on each day’s hand kinematics separately and computed the variance explained by each principal component.

#### Task feedback density

RP perturbations maintained the task plane aligned to the learned baseline mapping, whereas RA perturbations changed the task plane to a new subspace. A movement within the baseline *XY* plane therefore generates *cursor* movements with different amplitudes for each perturbation. Thus, the amount of task-relevant feedback can differ for the two perturbations. We investigated how this difference in feedback shaped learning by analyzing the relationship between hand and cursor movement variability. We defined feedback density as the ratio of cursor movement (confined to the 2D task plane) to total hand variance.

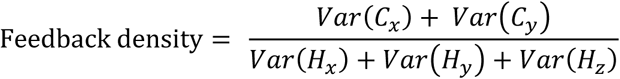

When feedback density is close to 1, all hand variability results in cursor movement. Cursor and hand variance were computed over the movement window (between the go cue and reaching the peripheral target) for each reach.

### Statistical analysis

Statistical significance was defined as *p* < 0.05 using statistical tests summarized below; details of test types, test direction, and sample sizes for each comparison are reported within the Results. Reported *p* values were mapped to significance levels using the following convention: p < 0.0001 (****), p < 0.001 (***), p < 0.01 (**), p < 0.05 (*), and p > 0.05 (n.s). Values are reported as mean ± standard deviation unless otherwise noted. All analyses were implemented in Python using the scipy.stats library. Repeated measures, such as success percentage, cursor error, and variance were compared across behavioral blocks using paired t-tests. We compared mean success percent during baseline mapping session A to early and late phases of the perturbation and washout block for statistical comparisons across series. One sample t-tests were used to assess whether within day learning, offline gains and performance changes between consecutive days were significantly above zero. Linear regression was used to evaluate relationships between perturbation magnitude and learning related measures including time to learn, washout magnitude and null axis variance. When quantifying across series effects, data from both monkeys were pooled and individual monkeys were treated as repeated instances of the same experimental design, which is standard practice in non-human primate studies.

## Results

### Animals identified task-relevant dimensions in baseline mapping

We first examined how monkeys learned cursor control in our redundant task, where their hand movements in 3D space controlled 2D cursor movements (Fig. 1A). Both monkeys initially practiced with a baseline mapping (*M_base_*), which translated horizontal and vertical hand movements to cursor position, while front– back hand motion had no effect on cursor position (Fig. 1B). After training, animals attained high task proficiency, evidenced by high success rates (Fig. 1C, top) and straight cursor trajectories (Fig. 1C, bottom) across 10 consecutive days (randomly selected from the period prior to beginning any perturbations).

The task allowed movement in three dimensions, but we found that the monkeys rapidly identified the lower-dimensional task-relevant space and restricted their movements primarily to a plane. Visual inspection of 3D hand trajectories showed that they lay almost entirely within a 2D subspace (Fig. 1D). Consistent with this, principal component analysis (PCA) of 3D reaches showed that the first two components captured over 95% of the movement variance (Fig. 1E). We used these top two PCs to capture the plane in which the animals primarily moved. We compared the variance captured by each animal’s movement plane (defined by their top two PCs) to that of the baseline task plane (*XY* plane) and found that points clustered near the unity line (Fig. 1F). We concluded that the animal’s learned movement plane was closely aligned to the task-relevant plane.

We also analyzed movements over the course of training to investigate how quickly animals identified their planar solution. Each animal performed thousands of reaching trials (Monkey A: 13,545; Monkey B: 48,342 trials) before perturbations were introduced, during which they rapidly reduced task-irrelevant movements. Their task-irrelevant variance had a half-life of only 122.4 trials for Monkey A and 693 trials for Monkey B and remained low thereafter (Fig. 1G). These results show the monkeys reduced task irrelevant movements early in their training, reflecting rapid learning of the planar task structure and accurate identification of the task-relevant plane.

### Relevance-preserving perturbations were learned within a training session and generalized

After training in the baseline mapping, we introduced different types of perturbations. Relevance-preserving (RP) visuomotor perturbations rotated the hand-to-cursor mapping but maintained the same task-relevant plane as the baseline mapping (see Methods; Fig. 2A). Introducing RP perturbations transiently disrupted task performance, causing an initial drop in success rate that rapidly recovered within the session (Fig. 2B, top, Fig. 2C; early B success on day 1 < baseline mean success, p = 0.007 (**); late B success on day 1 < baseline mean success, p = 0.19 (n.s.), one-sided paired t-test, N = 15 RP learning series across two monkeys). When the original baseline mapping was reinstated (A’ block), we observed a transient performance drop, which again resolved within the session (Fig. 2B top, Fig. 2C; early A’ success on day 1 < baseline, p = 0.00645 (**); late A’ success on day 1 < baseline, p = 0.186 (n.s.), one-sided paired t-test, N = 15 RP learning series across two monkeys).

Monkeys were exposed to the same RP perturbation across days within a learning series (9 of 15 series spanned multiple days), allowing us to examine learning dynamics across days. On re-exposure to the same perturbation on day 2, monkeys began at a slightly higher task performance compared to day 1 (Fig. 2B bottom, early B success percent: day 2 mean = 75.5 ± 16.6%, day 1 mean = 57.4 ± 20.6%, day 2 – day 1 success > 0, p = 0.0023 (**), one-sided one sample t-test, N = 9 RP learning series with multiple days). To capture learning within and across sessions, we quantified the speed of learning for each perturbation size by calculating the number of trials required to consistently reach 80% task success across all perturbation sessions in a learning series (time to learn; see Methods). Time to learn increased with perturbation magnitude (Fig. 2D, R² = 0.408, p = 0.00778 (**), linear regression), consistent with observations in many visuomotor perturbation experiments without control redundancy (Krakauer et al., 2000a; Marko et al., 2012).

We next analyzed the structure of cursor trajectories during learning. Introducing the RP perturbations resulted in curved trajectories early in learning, which gradually became straighter over the course of learning (Fig. 2E). Returning to baseline (A′ block) resulted in trajectories that curved in the direction opposite of the perturbation, consistent with generalization patterns commonly observed in visuomotor adaptation (Krakauer et al., 2019). We quantified this effect by calculating the deviation relative to a straight-line trajectory (cursor error, see Methods: Quantifying washout; example series shown in Fig. 2F). Cursor error increased when perturbations were introduced (first day: early cursor error in B > mean cursor error in A, p = 0.000095 (****), one-sided paired t-test, N = 15 RP learning series). Practice within a session during B led to a decrease in cursor error (first day: late cursor error in B < early cursor error in B, p = 0.00008 (****), one-sided paired t-test, N = 15 RP learning series). However, cursor error during the perturbation did not fully recover to baseline levels within a session (late cursor error in B > mean cursor error in A, p = 0.00039 (***), one-sided paired t-test, N = 15 RP learning series; mean absolute difference from baseline = 1.033, range 0.112–2.153). Cursor trajectories in A’ started with a higher cursor error compared to baseline across the learning series (first day: early cursor error in A’ > mean cursor error in A, p = 0.000113 (***), one-sided paired t-test, N = 15 RP learning series). These errors decreased within a session (first day: late cursor error in A’ < early cursor error in A’, p = 0.007 (**), one-sided paired t-test, N = 15 RP learning series). Moreover, cursor error reversed sign upon returning to baseline mapping (A’ block), similar to generalization observed in visuomotor adaptation (Krakauer et al., 2019). The magnitude of these generalization effects, assessed by average cursor error in the first 60 trials of washout, was correlated with the size of the preceding RP perturbation (Fig. 2G, R² = 0.8699, p = 0.0001 (***), linear regression).

We also analyzed cursor error patterns across repeated exposure to the same RP perturbations during learning series. Initial cursor error in the perturbation block decreased across days (early cursor error on first day > last day, p = 0.0159 (*), one-sided paired t-test, N = 9 RP learning series with multiple training days across 2 monkeys). However, it did not fully return to baseline levels (last day: early cursor error in B > mean cursor error in A, p = 0.00031 (***), one-sided paired t-test, N = 9 RP learning series). Washout of the learned adaptation in A’ was also observed across multiple days (early cursor error in A’ > mean cursor error in A, first day: p = 0.000036 (****), last day: p = 0.00007 (****), one-sided paired t-test, N=9). Thus, while across-day learning occurred, the majority of adaptation took place within individual sessions, and generalization patterns did not change with multi-day learning.

In a redundant task, RP perturbations may also affect behavior beyond the task-relevant plane, such as 3D hand movements. We investigated whether hand movements remained planar during RP perturbations and, if so, how the monkeys’ movement plane related to the task-relevant plane (the *XY* plane, as with the baseline mapping). Qualitatively, hand trajectories remained largely within a single plane during the RP perturbations and upon returning to baseline after exposure to the perturbation (Fig. 2H). We quantified planarity using the proportion of movement variance captured by the first two principal components of hand kinematics on the last session of each perturbation learning series when performance had plateaued and behavior was most stable. A large portion of variance was captured by the first two principal components (mean ± s.d.; Monkey A: 0.87 ± 0.06, N = 8 mappings; Monkey B: 0.93 ± 0.03, N = 7 mappings). We then computed the portion of movement variance captured by the task-relevant (XY) plane (mean ± s.d.; Monkey A: 0.94 ± 0.06, N = 8 mappings; Monkey B: 0.92 ± 0.05, N = 7 mappings). The proportion of movement variance within the two planes were similar (Fig. 2I; two-sided paired t-test, p = 0.148 (n.s), N = 15 mappings across both monkeys), indicating that hand movements remained planar and closely aligned within the task-relevant XY plane across adaptation to RP perturbations.

Despite its small relative contribution to overall movement, we examined null-axis (Z) variance to further analyze changes to task-irrelevant movements across learning. Variability along the null axis (Z) remained low across all blocks (comparison of mean null axis variance across the A, B and A’ blocks on day 1, F = 3.19, p = 0.051 (n.s), one-way ANOVA, N = 15 RP learning series) (Fig. 2J). Exposure to the perturbation did not significantly affect null axis movements early in the block (Day 1: early null variance in B vs. mean null variance in A, p = 0.356 (n.s), two-sided paired t-test, N = 15) nor late in the block (Day 1: late null variance in B vs. mean null variance in A, p = 0.643 (n.s), two-sided paired t-test, N = 15). Exposure to RP perturbations also did not alter task-irrelevant movements upon returning to the baseline block (A’) (Day 1: early null variance in A’ vs. mean null variance in A, p = 0.989 (n.s); late null variance in A’ vs. mean null variance in A, p = 0.986 (n.s), two-sided paired t-test, N = 15 RP learning series). Variability along the null axis during the first 60 trials of the washout block similarly did not show any relationship with perturbation size (Fig. 2K, R = -0.145, p = 0.637 (n.s), linear regression), confirming that generalization effects in washout were restricted to the task-relevant plane.

### Relevance-altering perturbations required multi-day learning and showed no generalization

We next examined learning in relevance-altering (RA) perturbations, which rotated the task-relevant plane into a new orientation in 3D hand space and therefore altered which dimensions of hand movement produced task-relevant feedback (Fig. 3A). Introducing RA perturbations substantially reduced the proportion of successful trials (Fig. 3B, top; early B success on day 1 < baseline mean success, p = 0.0014 (**), one-sided paired t-test, N = 7 across two monkeys). Unlike RP perturbations, monkeys were generally unable to recover baseline-level performance within a single session (Fig. 3B top, late B success on day 1 < baseline mean success, p = 0.0129 (*), one-sided paired t-test, N = 7 across two monkeys). However, performance gradually improved across days of exposure to the same RA perturbation (Fig. 3B, bottom, early B success on day 1 < early B success on day 2, p = 0.0037 (**); early B success on day 2 < early B success on last day, p = 0.0011 (**), one-sided paired t-test, N = 6 series with consecutive days) and eventually matched baseline performance (early B success on last day < baseline mean success, p = 0.0931 (n.s), one-sided paired t-test, N = 6), suggesting slow acquisition of a new control strategy. Success percent quickly returned to near-baseline levels when the baseline plane was reintroduced during A’ block. This rapid recall occurred both on the first day, where little to no learning occurred (Fig. 3C, early A’ success < baseline mean success, p = 0.262 (n.s), one-sided paired t-test, N = 7 RA learning series across two monkeys), as well as the last day when the RA mapping was well-learned (early A’ success < baseline mean success, p = 0.105 (n.s), one-sided paired t-test, N = 7). The time to learn RA perturbations did not show any consistent relationship with the perturbation size (Fig. 3D, R² = 0.1, slope = 1.96, p = 0.442 (n.s), linear regression), a notable contrast to RP perturbations (see Fig. 2D).

Unlike RP perturbations, where cursor error changes occurred largely within a single session, RA cursor error changes evolved across multiple days of training. Qualitatively, RA cursor trajectories did not show systematic error patterns during the perturbation (B block) nor a clear mirroring of errors upon return to baseline (A’ block) (Fig. 3E), a further contrast to the adaptation and generalization patterns seen in RP perturbations (Fig. 2E). The new RA mappings did consistently increase cursor errors compared to baseline (Fig. 3F, early cursor error in B vs. mean cursor error in A, first day: p = 0.00498 (**), last day: p = 0.0453 (*), two-sided paired t-test, N = 7 RA learning series). Within-day practice did not significantly decrease these errors (cursor error late < early on the first day with at least 20 trials, p = 0.0522 (n.s), paired t-test, N = 7 RA learning series). However, with extended practice, cursor error in B returned to baseline levels (late cursor error in B vs. mean cursor error in A, first day: p = 0.00762 (**), last day: p = 0.147 (n.s), two-sided paired t-test, N = 7 RA learning series). Even after RA perturbations were well-learned, cursor error patterns did not generalize between the perturbation and baseline mappings; cursor errors in the post-perturbation block (A’) were similar to those in baseline (early cursor error in A’ vs. mean cursor error in A, first day: p = 0.327 (n.s), last day: p = 0.1 (n.s), two-sided paired t-test, N = 7). The magnitude of washout during the A′ block, assessed by average cursor error in the first 60 trials, did not show any relationship with perturbation size (Fig. 3G, R = -0.286, p = 0.281 (n.s), linear regression). Thus, RA perturbations produced no generalization even when well-learned, in contrast to the adaptation-like washout in RP perturbations.

We next examined how 3D hand kinematics changed during RA perturbations. Hand kinematics during RA perturbations initially did not appear planar (Fig. 3H). However, movements became planar by the end of learning; the first two principal components of hand kinematics captured a large proportion of the total movement variance on the last day of each RA mapping (mean ± s.d.; Monkey A: 0.90 ± 0.02, N = 3 mappings; Monkey B: 0.92 ± 0.01, N = 4 mappings). We then measured the proportion of movement variance captured by the task relevant plane for each RA perturbation (mean ± s.d.; Monkey A: 0.78 ± 0.09, N = 3 mappings; Monkey B: 0.70 ± 0.10, N = 4 mappings). The proportion of movement variance in the task-relevant plane was significantly lower than that of the monkey’s top two PCs (paired t-test, p = 0.0043 (**), N = 7 mappings across both monkeys, Fig. 3I). Thus, the monkeys generated largely planar movements to solve RA perturbations, but their movement planes did not consistently align with the task-relevant plane. This contrasts with RP perturbations, where hand movements remained planar and aligned with the task-relevant plane (Fig. 2I).

Upon returning to the baseline mapping, 3D hand kinematics resembled those of baseline (Fig. 3H), consistent with the lack of generalization across blocks seen in cursor movements. We assessed generalization by quantifying hand movement along the axis of movements that was task-null in the baseline condition (the Z-axis, orthogonal to the baseline plane). As expected, hand movement variance in the Z-axis during the perturbation block increased relative to that of baseline as animals learned the RA mapping (Fig. 3J, early null axis variance in B > mean null axis variance in A, first day: p = 0.0896 (n.s), last day: p = 0.00413 (**); late null axis variance in B > mean null axis variance in A, first day: p = 0.0139 (*), last day: p = 0.0296 (*), one-sided paired t-test, N = 7). Yet, animals did not appear to generalize changes in 3D hand movements to the baseline mapping. Task-irrelevant Z-axis movement variance returned to similar levels as that of baseline when the baseline mapping was reintroduced (Fig. 3J, early null axis variance in A’ > mean null axis variance in A, first day: p = 0.139 (n.s), last day: p = 0.629 (n.s) ; late null axis variance in A’ > mean null axis variance in A, first day: p = 0.154 (n.s), last day: p = 0.0771 (n.s), one-sided paired t-test, N = 7). Hand movement variance along the Z axis during the first 60 trials of the washout block did not show any relationship with the perturbation size (Fig. 3K, R = -0.139, p = 0.792 (n.s), linear regression). Thus, movements generated to explore the 3D space during RA learning did not carry over to the baseline mapping, regardless of perturbation size.

### Distinct learning timescales in RP vs RA perturbations

The prior analyses revealed that across-day learning patterns differed between relevance preserving (RP) and altering (RA) perturbations. We investigated the learning timescales for each perturbation type by comparing performance changes within and between sessions (Fig. 4). These two perturbations impacted performance differently (first day success rates for RA < RP mappings, p = 0.00223 (**), one-sided Mann-Whitney U test, N = 15 RP learning series, 7 RA learning series across both monkeys), which may provide an intuitive explanation for why RA perturbations took much longer to learn. However, *when* performance improved notably differed as well, as illustrated in representative sessions from RP (Fig. 4A) and RA (Fig. 4B) perturbations. RP perturbations produced rapid adaptation within each block, with animals reaching near-baseline success rates within a day and retaining only modest learning across days. In contrast, RA perturbations exhibited slower within-day gains and large performance jumps between consecutive days (offline gains). Motor learning computations that occur during task performance are thought to be both behaviorally and neurologically distinct from those that occur after performing the task (Dayan and Cohen, 2011; Krakauer et al., 2019). This observation, then, suggests that introducing large task errors by changing task-relevant spaces may alter the learning computations that are engaged.

We studied this by comparing within-session and across-session learning between perturbation types. Both RP and RA perturbations showed performance improvements within a single training session (Fig. 4C, mean within session performance change > 0, RP: 20.66% ± 15.9%, p = 1.65e-10 (****), N = 41 learning days across 15 RP learning series from two monkeys; RA: 21.42% ± 16.53%, p = 6.87e-8 (****), N = 29 learning days across 7 RA learning series from two monkeys; one-sided one sample t-tests). In contrast, RP and RA perturbations showed distinct patterns of offline gains (Fig. 4D). Only RA perturbations showed statistically significant offline performance changes (mean offline gain in performance > 0, RP: 2.85% ± 26.70%, p = 0.302 (n.s), N = 24 day pairs across 9 multi-day RP learning series from two monkeys; RA: 12.26% ± 12.15%, p = 8.145e-5 (****), N = 21 day pairs across 7 RA learning series from two monkeys; one-sided one sample t-tests). Across all perturbation training sessions, regardless of perturbation type, offline gains were inversely correlated to within-day behavioral improvements (R = -0.50, p = 0.011 (*); Fig. 4E), suggesting that offline learning primarily occurred after sessions where little learning occurred during practice.

We next asked what factors might influence learning dynamics across different perturbations. Motivated by the fact that RP and RA perturbations define different task-relevant spaces, we hypothesized that the amount of offline learning might be influenced by the extent to which the movement variance structure animals adopted in the baseline mapping produced task feedback. Animals generated movements that were primarily within the learned 2D baseline plane during RP perturbations (Fig. 2). Since the task-relevant plane is shared between the baseline and RP mappings, most of the animals’ movement variance resulted in informative cursor feedback. RA perturbations, by necessity, required animals to generate hand movements outside of the baseline plane to successfully perform the task (Fig. 3). However, we found that animals continued to generate movements in the baseline task-relevant plane during early exposure to the RA perturbations (Fig. 3H, 4F), which yielded little informative cursor feedback because it is a task-irrelevant subspace for the RA mapping. Example hand movement and cursor trajectories from early in the first RA and RP sessions illustrate this difference (Fig. 4F). We quantified this phenomenon by computing the ratio of cursor movement variance to hand movement variance, which captures how much of the animals’ movements contribute to visual task feedback (*Feedback density*, see Methods). Sessions with low feedback density were associated with larger offline gains (Fig. 4G, R2 = -0.29, p = 0.0037 (**), linear regression), indicating that the availability of task-relevant feedback may be one factor that contributes to whether learning occurs within a session or offline.

### History dependent modulation of movement variability

Feedback density depends on the geometric relationship between the task-relevant space and movement variability. Yet the time to learn RA perturbations did not correlate with perturbation size (Fig. 3D), indicating that perturbation magnitude and availability of task feedback alone were insufficient to explain learning dynamics when task relevance changed. We hypothesized that animals learned a shared structure underlying RA perturbations – information about the space of possible task-relevant planes – and used that learning to shape their behavior. This hypothesis predicts that prior experience with RA perturbations should have two consequences: (1) faster learning of subsequent RA mappings, and (2) changes to how animals generate movements to learn new RA mappings.

We tested whether experience with RA perturbations reduced learning time for subsequent RA perturbations. Time to learn RA perturbations decreased with training order irrespective of perturbation size (Fig. 5A, left). This observation was particularly striking when comparing the training trajectories of each monkey. Monkey A first experienced a small (20°) RA perturbation while Monkey B experienced a large (90°) one, yet both animals took the longest time to learn the first RA mapping they were exposed to compared to others (Fig. 5A, left). No such trend was observed for relevance-preserving (RP) perturbations (Fig. 5A, right), where time to learn was well-explained by perturbation size (see Fig. 2D). This history-dependent learning isolated to one type of perturbation is consistent with our hypothesis that structure learning contributed only to RA perturbation learning.

**Figure 5:**
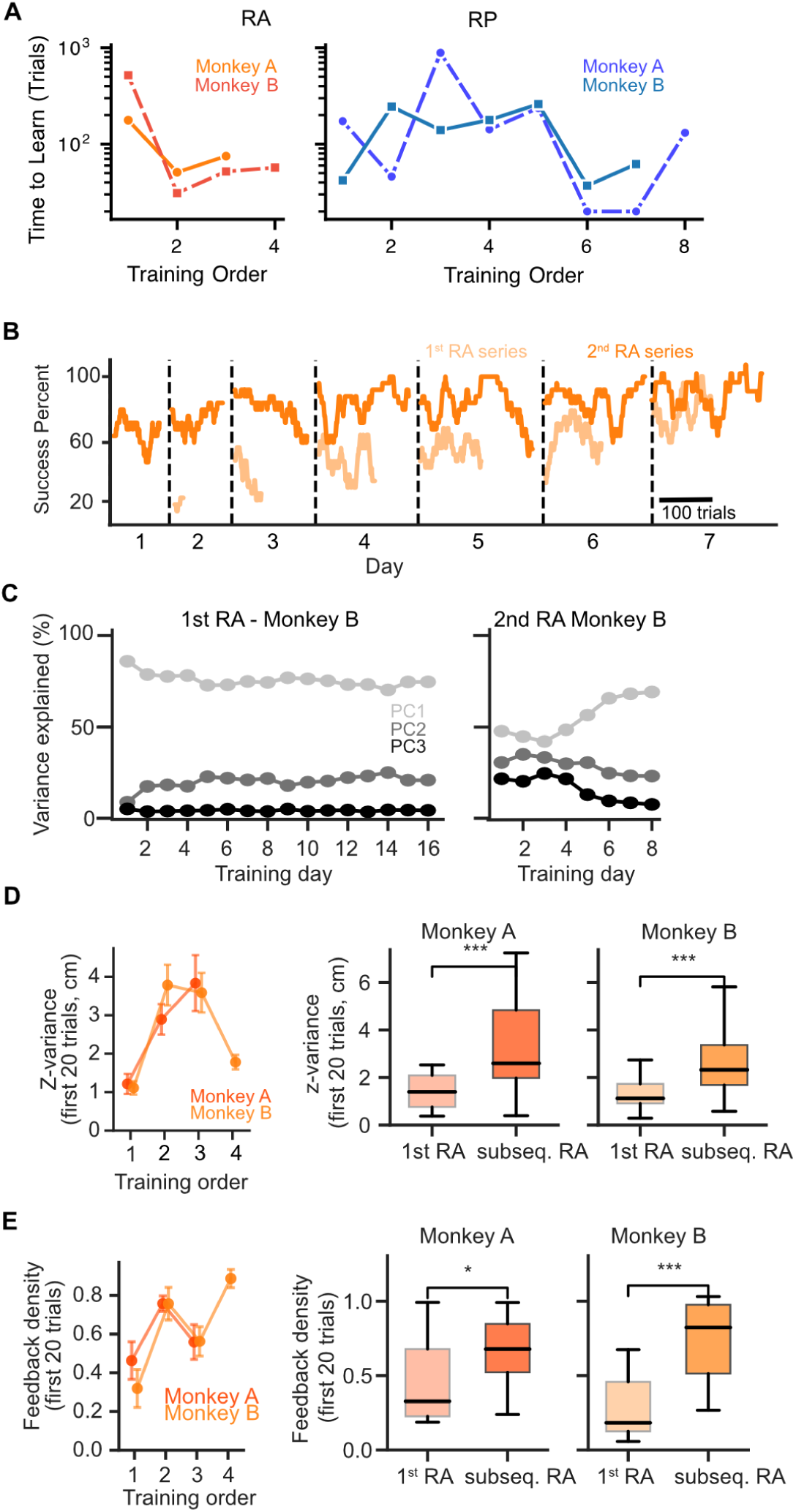
Prior learning influences exploration. **A)** Time to learn (log scale) as a function of training order for relevance-altering (RA; left, orange) and relevance-preserving (RP; right, blue) perturbations for Monkey A (solid lines) and Monkey B (dashed lines). **B)** Success percent for RA perturbation blocks across trials in two RA learning series of the same magnitude experienced at different times (first series, light orange; second series, dark orange) in Monkey B’s training history. **C)** Proportion of hand movement variance explained by the first (light gray), second (dark gray), and third (black) PC for first (left) and second (right) RA perturbation learned by Monkey B. **D)** Left: Mean variance along the Z axis (null in baseline) from the first 20 perturbation trials of each RA learning series as a function of training order. Error bars indicate standard error of the mean. Right: Mean Z variance in the first RA mapping compared to all subsequent RA mappings for two monkeys (right). Boxes indicate inter-quartile range, lines indicate medians, whiskers extend to 1.5x IQR. (Mean Z-variance from first 20 trials subsequent RA mappings > first RA mapping, Monkey A: p = 0.00031 (***); Monkey B: p = 0.000032 (****); Mann-Whitney U test). **E)** Left: Mean feedback density from the first 20 perturbation trials of each RA learning series as a function of training order. Error bars indicate standard error of the mean. Right: Feedback density in first RA mapping compared to subsequent RA mappings for two monkeys. Boxes indicate inter-quartile range, lines indicate medians, whiskers extend to 1.5x IQR. Feedback density from first 20 trials subsequent RA mappings > first RA mapping, Monkey A: p = 0.0495 (*); Monkey B: p = 0.00076 (***); Mann-Whitney U test.

If structure learning underlies this history dependence, prior experience would also alter how animals generate movements to learn new mappings. We first examined how one animal generated movements when they learned two closely related RA mappings, one for the first time and another after experience with RA mappings. During the first RA exposure, which used an *XZ* task-relevant plane (right–left hand movements controlled horizontal cursor motion and front–back movements controlled vertical motion), Monkey B exhibited slow, multi-day learning (7 days to exceed 80% correct) with little within-session improvement during early days. When Monkey B was later exposed to another RA perturbation with the same magnitude (same task-relevant plane, but a different within-plane mapping to cursor movement), he achieved higher initial success from the first day and improved performance in the RA blocks notably faster (2 days to exceed 80% correct) (Fig. 5B). The monkey’s 3D hand movements confirmed that he generated movements differently as he learned these two RA mappings. During his first exposure, Monkey B continued to make primarily planar, 2D reaches within the 3D workspace (Fig. 5C, left). During the second exposure, he immediately generated movements that spanned the 3D space before gradually converging to a stable 2D subspace over the course of several days (Fig. 5C, right). This example raises the possibility that structure learning, over the course of RA perturbation exposures, informed *exploration strategies* to identify changes in task-relevant spaces.

We analyzed movement variance patterns to confirm that this change in exploration generalized across all RA exposures. Because the Z-axis was task-irrelevant under the baseline mapping and became task-relevant under RA perturbations, movement variance along this axis provides an index for how much animals explored newly task-relevant dimensions. We measured hand movement variance along the Z-axis during the first 20 trials of each RA exposure, irrespective of task success, before task performance improved. Both monkeys generated more Z-axis variance during subsequent RA mappings than during their first exposure (Fig. 5D, mean Z-variance from first 20 trials subsequent RA mappings > first RA mapping, Monkey A: p = 0.00031 (***); Monkey B: p = 0.000032 (****); one-sided Mann-Whitney U test). This held irrespective of the magnitude of the subsequent perturbation, mirroring the size-independent reduction in time to learn (Fig. 5A).

If monkeys indeed learned useful information about the structure of RA mappings, we would predict that they weren’t only exploring *more*, but in ways that increased their ability to identify the task relevant plane. We began investigating this hypothesis by asking if monkeys learned to produce movements that generated more visual feedback during early trials. We returned to our feedback density metric to capture how the animals’ movement variability was oriented relative to the task-relevant plane, focusing specifically on the first 20 attempted trials. Both animals generated movements that produced more cursor movement (higher feedback density) on subsequent RA exposures than on the first (Fig. 5E, feedback density from first 20 trials subsequent RA mappings > first RA mapping, Monkey A: p = 0.0495 (*); Monkey B: p = 0.00076 (***); one-sided Mann-Whitney U test). Prior RA experience was therefore accompanied by both broader exploration of movements within the redundant environment and more task-relevant feedback, though we cannot fully separate the causal relationships or underlying mechanisms.

## Discussion

Adapting to sensory-motor maps that change the task-relevant space (RA perturbations) involved learning mechanisms beyond those used to adapt to new mappings within a learned task-relevant space (RP perturbations). Monkeys rapidly adapted to RP perturbations and their learning patterns were consistent with error-based adaptation. Monkeys adapted to RA perturbations slowly and their learning lacked hallmarks of error-based adaptation, instead showing patterns consistent with contributions from structure learning. Our task revealed these distinctions by isolating changes in task-relevant spaces from other factors like task goals or biological constraints. Our results highlight the importance of task-relevance learning computations in redundant tasks and provide a new paradigm to study many forms of motor learning.

### Initial task-relevance learning

Our redundant motor task allowed monkeys to move in 3D, but they rapidly constrained their movements to a lower dimensional subspace in their initial training (Fig. 1E). Their learned movements closely aligned with the task-relevant space (Fig. 1F, G), suggesting animals identified the overall *structure* of the task’s movement-outcome associations. Changes in task-irrelevant variability during initial learning may shed light on competing motor control hypotheses. The Uncontrolled Manifold (UCM) hypothesis posits that movements are not altered unless they impact task outcomes (Scholz and Schöner, 1999; Müller and Sternad, 2004), while the Optimal Feedback Control (OFC) hypothesis predicts that learning alters any property of movement that influences the system’s cost function (Todorov and Jordan, 2002). Rapid suppression of task-irrelevant variability is more consistent with the latter view, suggesting that the motor system likely optimizes factors like metabolic cost. While our study does not definitively resolve these theories, our task provides an experimental method to test predictions about how control policies shape movement and learning.

### Changing task relevance evokes distinct learning regimes

Selectively perturbing task-relevance revealed that the relationship between a previously-learned and new task-relevant space influenced learning dynamics (Figs. 2 and 3). Our results suggest this is due, in part, to differences in the feedback available to guide learning (Fig. 4F). Even if RP perturbations reduce task performance, they provide visual feedback about movement-cursor relationships. This corresponded with learning that had clear signatures of non-redundant visuomotor adaptation, which is dominated by error-based computations to update an internal model of movement-cursor relationships (Krakauer et al., 2000b, 2005; Krakauer, 2009; Shadmehr et al., 2010). In contrast, RA perturbations reduce task performance but can also reduce the amount of visual feedback about movement-cursor relationships. Indeed, for some geometric relationships, movements within a previously learned task space could produce *no* cursor movement, likely reducing the ability to perform error-based learning. Consistent with this, learning RA perturbations lacked signatures of visuomotor adaptation, and instead showed several similarities to *de novo* skill learning (Telgen et al., 2014; Yang et al., 2021). Moreover, RA perturbations produced larger changes in movement variability (Fig. 2I vs. 3I), which are thought to underlie exploration during skill acquisition (Dhawale et al., 2017). Together, these results suggest that adapting to new task-relevant spaces involves more than error-based adaptation, likely engaging mechanisms involved in skill learning (discussed further below).

Differences in feedback may also contribute to the distinct timescale for learning RA and RP perturbations. RP performance improved primarily within sessions, consistent with online mechanisms that dominate error-based adaptation (Smith et al., 2006). In contrast, RA performance often improved between sessions, consistent with offline learning in motor skill acquisition (Dayan and Cohen, 2011; Bönstrup et al., 2019; Krakauer et al., 2019). Regardless of perturbation type, sessions where animals experienced less visual feedback of their hand movements were followed by larger offline gains (Fig. 4G). This observation generates the hypothesis that learning shifts to offline mechanisms when sensory feedback of task errors, which are known to heavily drive online learning (Shadmehr et al., 2010), is limited. Future experiments pairing our redundant perturbation paradigm with neural measurements will provide ways to test this hypothesis and study the neural mechanisms governing offline and online learning.

### Structure learning, context, and history dependence

Longitudinal training revealed further distinctions in the timescales of learning for RP and RA perturbations. The time required for animals to learn RA perturbations was not solely determined by perturbation size, and depended on their history of exposure to this type of perturbation (Fig. 5A). RP perturbations did not show a history dependence, providing further evidence that the two perturbations engage distinct learning mechanisms. Meta-learning is also observed in models that incorporate context-dependence (Heald et al., 2021) and in skilled behaviors that require structure learning (Kobak and Mehring, 2012), suggesting these computations may uniquely contribute when adapting to changes in task-relevance. Consistent with this, animals changed their movement variability patterns after exposure to different task-relevant spaces, likely based on learned or inferred task structure (Fig. 5C-E). Observations from skill learning, primarily in songbirds and rodents, show that the nervous system modulates variability within motor circuits to leverage its benefits for skill acquisition while minimizing disruptions to precise execution (Drew and Abbott, 2003). Our work suggests that context and structure inferences – computations often thought of as cognitive and primarily probed in human behavior – contribute to generating motor variability. Task-relevance perturbations open new opportunities to investigate the neural circuits involved in generating motor variability and how these are informed by cognitive computations.

The influence of training history within our task introduces richness that will improve our understanding of motor learning but also highlights new experimental considerations. For instance, our results suggest that RA perturbation learning was also influenced by their sequencing. Our randomized design resulted in each monkey experiencing different initial RA perturbations – small in Monkey A, large in Monkey B. They, correspondingly, appeared to change their exploration strategies in different ways across RA perturbations – gradually in Monkey A, rapidly in Monkey B (Fig. 5D-E) – but the cohort constraints of monkey studies limit our ability to prove this relationship. Conversely, animal models allowed us to precisely control each subject’s prior task familiarity and training history, which is infeasible in human subjects who perform closely related behaviors in daily life (Turnham et al., 2011). Monkeys provide precise ways to probe initial task learning from a naïve state (Fig. 1) and all subsequent exposures. Examining meta-learning and its neurophysiological basis will, therefore, likely benefit from approaches that combine the statistical power of high throughput behavioral studies (in humans and/or animals) and neurophysiology experiments.

### Implications for Models and Neural Manifolds

Our results reinforce that motor learning can span timescales from months-long skill learning to single-trial recall (Abbott and Dayan, 2005) and models suggesting that these arise through a combination of more “cognitive” processes like context inference paired with “proper” sensorimotor learning (Heald et al., 2021). In that framework, our results suggest that RA and baseline mappings were treated as distinct contexts since there was little cross-task generalization, and that the long time-scales of initial RA learning were due to it being a novel context (Fig. 3). The computational underpinnings of the history-dependence for RA perturbations, however, may require modeling advances. Few motor learning models address how movements are generated to learn within a context and largely assume action-outcome structures are known or observable. The history-dependence we observed in RA perturbations suggests that generating movements to learn action-outcome relationships was informed by the inferred context. Modeling interactions between “cognitive” and “motor” computations will likely be critical for understanding skill acquisition and adaptation.

Our results also add nuanced interpretations of studies probing the neurophysiological bases of skill acquisition. The slower timescales associated with learning new motor skills have been attributed to biological constraints like synaptic connectivity changes or muscle coordination patterns (Abbott and Dayan, 2005; d’Avella and Bizzi, 2005; Sadtler et al., 2014; Oby et al., 2019; Busch et al., 2025). Key evidence for this hypothesis comes from brain-computer interfaces, where mappings requiring neural activity “outside” the repertoire of patterns seen in the task (the “neural manifold”) are learned more slowly than mappings within that repertoire (Sadtler et al., 2014; Oby et al., 2019). Our work shows that a similar separation of timescales can emerge even when the movements a mapping requires are already within the motor system’s repertoire. Our monkeys presumably possessed the ability to generate the arm movements required for RA perturbations, but needed extensive practice to associate those movements with the new task context. Our results suggest that the timescale of learning may reflect the computations needed to infer which movements are task-relevant, alongside biological constraints. This interpretation, paired with our results linking limited feedback with structure learning (Fig. 4F, 5E), also aligns with the observation that *incomplete* feedback alone can reproduce brain-computer interface learning dynamics (Feulner and Clopath, 2021). Our task paradigm is well positioned to test how learning new task-relevant spaces engages synaptic plasticity and shapes the neural repertoire.

Our study demonstrates that redundant action-outcome relationships evoke a continuum of learning dynamics, from rapid adaptation to slow, history dependent learning, likely reflecting multiple interacting computations. Precise sensorimotor manipulations that consider task-spaces within redundant environments can evoke skill acquisition, adaptation, and structure learning, opening new opportunities to investigate their underlying neurophysiological mechanisms.

## Acknowledgments

We would like to thank the Washington National Biomedical Research Behavioral Management team for their consultations on animal training. This work was supported in part by an NSF Accelnet INBIC fellowship (P.R.), a postdoctoral fellow award from Weill Neurohub (L.R.S.), a Simons Collaboration for the Global Brain Pilot award (898220, A.L.O.), the National Institute of Neurological Disorders and Stroke (NIH R01 NS134634, A.L.O.), and the NIH Office of Research Infrastructure Programs (P51 OD010425). We acknowledge Mathew Sarti’s work that explored change in variance as monkeys learned the baseline mapping.

